# Boundary-Driven Emergent Spatiotemporal Order in Growing Microbial Colonies

**DOI:** 10.1101/328583

**Authors:** Bhargav R. Karamched, William Ott, Ilya Timofeyev, Razan N. Alnahhas, Matthew R. Bennett, Krešimir Josić

## Abstract

We introduce a tractable stochastic spatial Moran model to explain experimentally-observed patterns of rod-shaped bacteria growing in rectangular microfluidic traps. Our model shows that spatial patterns can arise as a result of a tug-of-war between boundary effects and modulations of growth rate due to cell-cell interactions. Cells align *parallel* to the long side of the trap when boundary effects dominate. However, when the magnitude of cell-cell interactions exceeds a critical value, cells align orthogonally to the trap’s long side. Our model is analytically tractable, and completely solvable under a mean-field approximation. This allows us to elucidate the mechanisms that govern the formation of population-level patterns. The model can be easily extended to examine various types of interactions that can shape the collective behavior in bacterial populations.

---

Patterns emerge in collectives of interacting biological agents even in the absence of leaders or global signals. Collective motions of birds and fish arise from simple interactions between neighbors [1, 2], gliding *M. xanthus* form coherently moving clusters via steric interference [3], and molecular motors self-organize to transport intracellular cargo [4]. Yet how the actions of myopic agents drive collective behavior is not fully understood.

Physical interactions between neighbors in microbial colonies also lead to emergent patterns [5, 6]. For instance, multi-strain consortia of *E. coli* in open, rectangular microfluidic traps form single-strain bands (see Fig. 1a). Such spatial arrangements are stable and help maintain consistent dynamics in bacterial strains interacting via extracellular signals [7, 8]. Uncovering how local interactions drive population-level patterns is thus important for understanding emergent order in bacterial communities and engineering synthetic microbial collectives with desired properties [9–11].

**FIG. 1.**
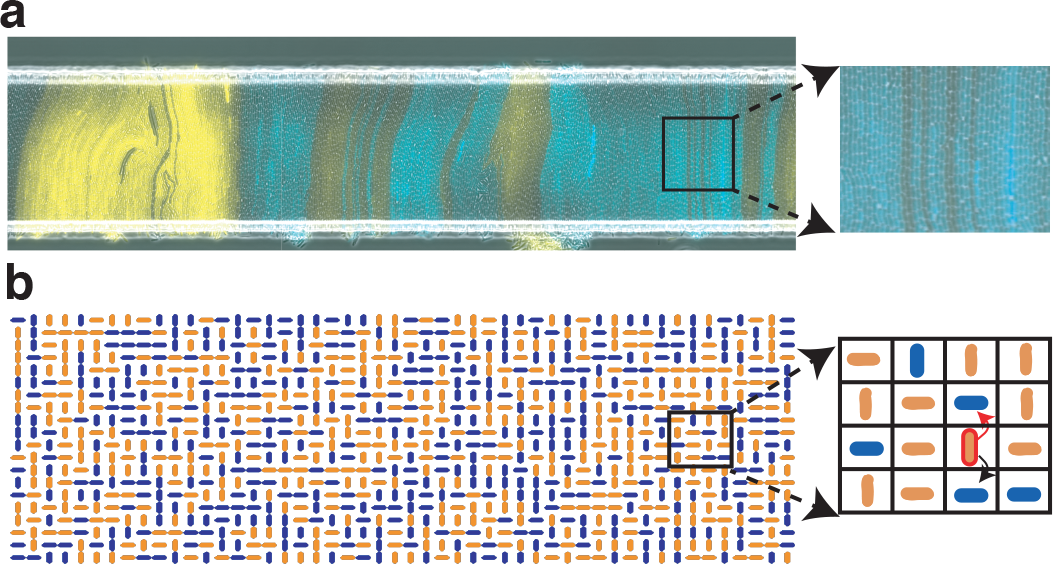
(Color online) (a) A monolayer of *E. Coli* in an open microfluidic trap with cells aligned orthogonally to the trap’s long side. Colors represent distinct strains. (b) In our spatial Moran model cell growth is directional and location dependent: The outlined vertical cell can grow only upward or downward at a location-dependent rate. The red arrow indicates growth direction, so the cell above will be replaced by a descendant of the outlined cell. We model single strain populations, but use the same color for mother and daughter cells for visualization.

Recent experiments and agent-based simulations suggest that environmental geometry and physical interactions between microbes influence observed global structures [12–15]. Capsule-shaped *E. coli* grow along the major axis of their bodies, preferring directions with minimal physical resistance, *i.e.* where the number of cells requiring displacement is smallest [6, 12, 16]. Moreover, in dense populations, cell growth is also determined by the geometry of the confining space [12, 14].

We provide a tractable mathematical model that shows how such directional, cell-interaction-dependent growth drives population-level patterns. We model cells as horizontally- or vertically-oriented agents on a lattice representing a microfluidic trap (see Fig. 1b). A cell’s orientation determines the directions in which it divides, while its location determines its growth rate. As physical growth requires the displacement of fewer cells towards the nearer boundary, we assume that division along this direction is more probable.

Our model shows that a transition occurs at a critical value of cell-cell interactions: When cells do not strongly impact each other’s growth, the collective aligns into columns *parallel* to the long side of a trap (see Fig. 2b). However, if the strength of these interactions is sufficiently strong, the collective aligns *orthogonally* to the trap’s long side (see Figs. 2a and 1a). Since the latter arrangement is observed experimentally, our model suggests that cell-cell interactions modulate growth rates [5, 14], and drive the emergence of ordered states in spatially-extended populations[17].

**FIG. 2.**
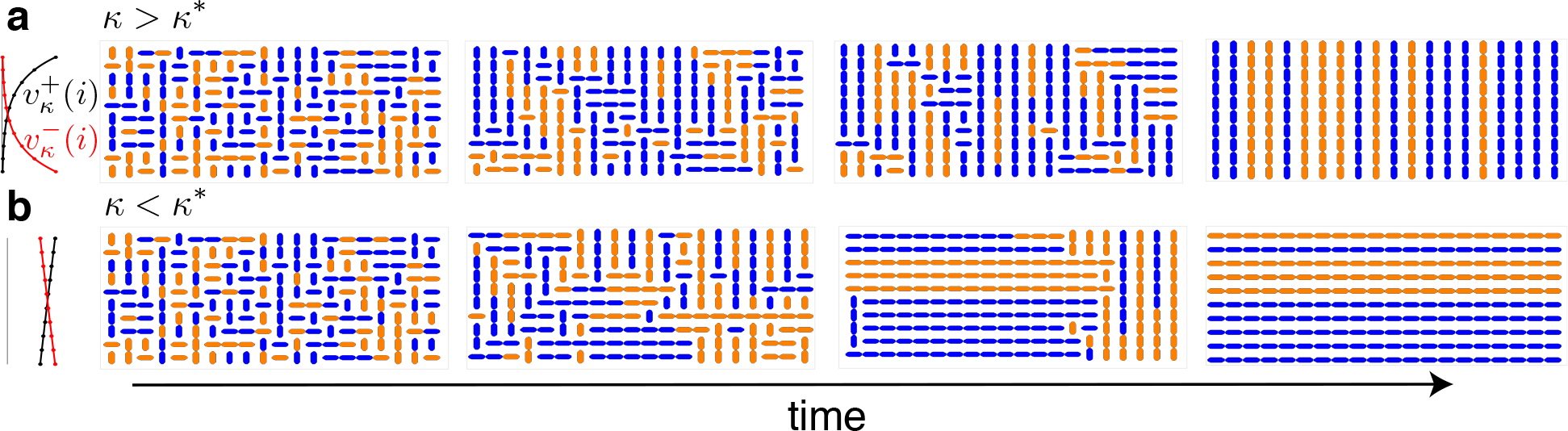
(Color online) Cells growing on a lattice according to a SMM. (a) Snapshots of the transient states and the all-vertical equilibrium for for *k* > *k*\*. On the left, we show growth rates of vertically-oriented cells toward the upper and lower boundaries; (b) Same as (a) but for *k* < *k*\*. See SI for corresponding movies.

Previous detailed models of growing bacterial colonies provided limited insight into the mechanisms underlying emergent structures. A growing bacterial population can be modeled as an expanding fluid with a dynamical order parameter reflecting cell alignment [13, 18–20]. Agent-based models can resolve physical interactions between cells in dense populations [15, 21, 22]. Both approaches can reproduce spatiotemporal patterns observed in cell collectives, but are difficult to analyze. We provide a simple, analytically tractable, and flexible alternative which offers insights into the emergence of spatial structures.

## Spatial Moran model (SMM)

Our model captures essential features of a population of rod-shaped bacteria growing in a microfluidic trap: To grow in the direction of the longer axis of their capsule-shaped bodies, cells displace their neighbors. Division results in a daughter cell approximately aligned with its parent. A cell at the trap’s boundary can be pushed out by cells growing in the interior, and bacteria at the boundary can produce offspring outside the trap.

We model the rectangular microfluidic trap as an *M* × *N* lattice filled by vertically- or horizontally-oriented cells (see Figure 1b). Initially the lattice is full, and cell orientation is random [23]. Cells grow at location-dependent rates. Upon division, a cell’s offspring replaces one of its neighbors. We denote by 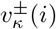 the growth rate of a vertical cell in the *i*th row toward the top (+) or bottom (−) boundary, and 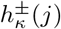 the growth rate of a horizontal cell in the *j*th column toward the right (+) or left (−) boundary (see Fig. 1b). The growth rate of a vertical (horizontal) cell depends only on the row (column) in which it resides.

We assume growth rates are determined by a one-parameter function family, with *κ* ∈ [0, ∞) characterizing the population’s impact on growth. This family can be general, but we assume that growth rates are positive and satisfy: (1) There exists a λ ∈ (0, ∞) such that 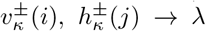 as *k* → 0 for all *i*,*j*; (2) Maximal growth rates occur at the boundaries, 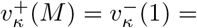 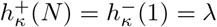; (3) 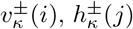 decrease monotonically with distance from the boundary that maximizes their value. Condition (1) says that cells grow uniformly at rate λ in the absence of interactions (*κ* = 0). Conditions (2) and (3) reflect cells’ tendency to grow toward the nearest boundary, and growth rate dampening from cells obstructing growth in a certain direction (see Fig. 1b). Unless otherwise noted, we used [24]

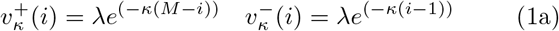

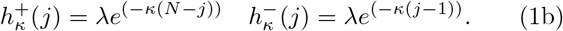

Cells grow by displacing their neighbors: In a small interval, Δ*t*, a vertical (horizontal) cell at the *ij*-th site replaces a neighbor at (*i* ± 1)*j* (respectively *i*(*j* ± 1)) with a copy of itself with probability 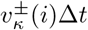 (respectively 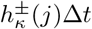). Divisions are independent across the population, and thus exponentially distributed. Only the division of an adjacent cell with the opposite orientation can alter the orientation at the *ij*-th site. Boundaries are absorbing, and divisions at the boundary producing descendants outside the trap result in no changes. In the microfluidic trap, cell growth and division physically displace more than one cell in the direction of growth. A model that incorporates such long-range interactions displays similar behavior (see SI Fig. S6 [25]).

## Results

To understand the impact of trap geometry on collective bacterial cell alignment, we simulated the SMM using the Gillespie algorithm [26] on lattices with different aspect ratios, Γ ≡ *N*/*M*, and different interaction parameters, *κ* [27]. For *κ* sufficiently large, all initial conditions converge to the equilibrium where cells are orthogonal to the long side of the trap (see Fig. S2; for Γ > 1, all cells vertical, for Γ < 1, all cells horizontal). When Γ = 1, the system reaches a quasiequilibrium with cells orthogonal to the nearest boundary (see Fig. 3). This suggests that Γ acts as a parameter for a transcritical-like bifurcation at Γ = 1 where the horizontal and vertical equilibria exchange stability. We make this precise in the next section.

**FIG. 3.**
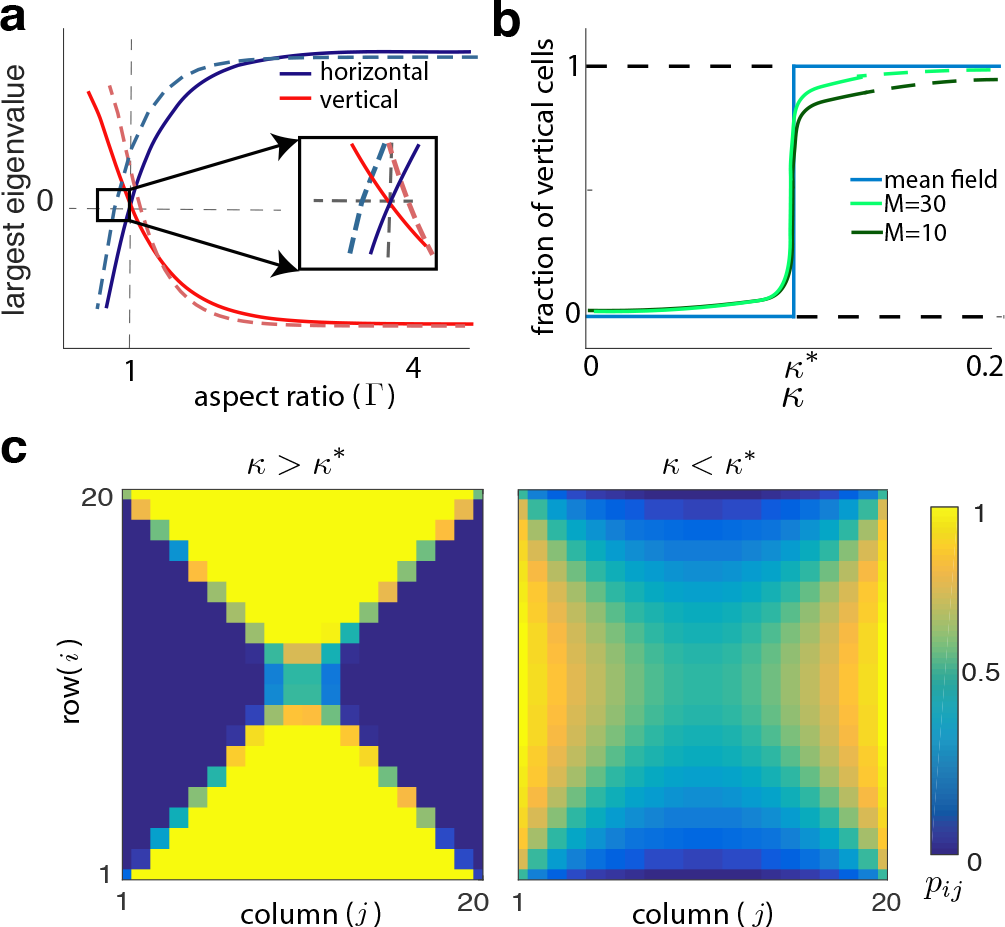
(Color online) (a) Eigenvalue plots as described in text for different system sizes (*M* = 10, dashed, *M* = 100, solid). The two states lose stability at different points for small systems, so that for a range of r neither is stable. In the large system the equilibria lose stability nearly simultaneously at Γ = 1. (b) The fraction of vertical cells at equilibrium exhibits a sharp transition near *k*\* for fixed Γ > 1 (Eq. (3), blue, and closed ME, Eq. (2), green). A secondary bifurcation in the all-vertical state occurs at *k* > *k*\* (solid to dashed green line transition) (c) Steady states of the closed ME when Γ = 1 for *k* > *k*\* and *k* < *k*\*.

Interestingly, when *κ* = 0, cells orient *parallel* to the long side of the trap (see Fig. 2b): When Γ > 1 (Γ < 1), the horizontal (vertical) equilibrium is stable. When Γ = 1, symmetry again results in a saddle-like quasiequilibrium, with cells *parallel* to the nearest boundary.

Therefore, when cells divide at location-independent rates, (*κ* = 0) they approach an equilibrium opposite to that when growth is location-dependent (*κ* is large). We observed the second state experimentally, suggesting that such cell-cell interactions influence global structure. The model also suggests that a phase transition occurs at a critical value, *k*\*.

This exchange of stability between equilibria at *k*\* results from an interplay between boundary effects and growth rate variations. When *κ* = 0, all cells divide at equal rates, except for those orthogonal to a boundary which are as likely to have a descendant within the trap as outside. However, more cells are likely to be orthogonal to the *long* boundary than the short one and to have a descendant outside the trap, so that cells *parallel* to the long boundary have a higher effective growth rate, and eventually fill the trap (see Fig. 2b). Conversely, when *k* > *k*\*, cells parallel to the longer side of the trap will have more cells obstructing them than cells parallel to the short side. If *κ* is sufficiently large, the average growth rate of cells perpendicular to the long boundary will dominate, and these cells will fill the trap (see Fig. 2a). Even when *k* > *k*\*, variations in growth rates across the lattice can be small: In a 20 × 10 lattice, *k*\* ~ 10^−2^ (see below) and cell growth is reduced by half at ≈ 70 cell lengths.

Cell-cell interaction kernels satisfying conditions (1)-(3) will generally lead to the same qualitative results, and we obtain the critical values *k*\* analytically for a range of different functions below. As expected, *k*\* → ∞ as lattice size grows, and near critical values in larger traps growth rates have smaller spatial variations than in smaller traps.

## Master equation model

To understand the dynamics of the SMM we develop a master equation (ME) describing the evolution of occupation probabilities at different lattice sites. Denote by *n*_*ij*_ ∈ {0,1} the state of the *ij*-th site at time *t*, so that *n*_*ij*_ = 1 (*n*_*ij*_ = 0) corresponds to a site occupied by a vertical (horizontal) cell. The dynamics of the probability *p*_*ij*_(*t*) of *n*_*ij*_ = 1 at time *t* are characterized by the ME [28, 29],

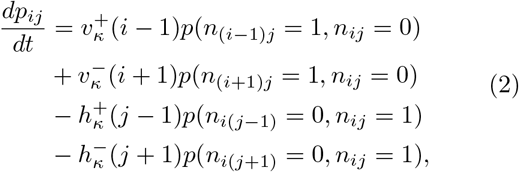

where *p*(*n*_*ij*_, *n*_*kl*_) are joint occupation probabilities. The first two terms in Eq. (2) correspond to horizontal-to-vertical cell transitions through displacement by a descendant from a cell either above or below. The second two terms describe the opposite transition. Equations at boundary sites are obtained by setting to zero the probability that a site outside the lattice is occupied, *e.g. p*(*n*_*i*_(*N*+1) = 0, *n*_*ij*_ = 1) = *p*(*n*_0*j*_ = 1, *n*_*ij*_ = 0) = 0.

Eq. (2) is related to the Ising model as both describe the evolution of alignment probabilities on a lattice. However, the location-dependent growth rates lead to different interactions, and no external field influences cell alignment [29] in our model.

This ME describes a nontrivial many-body problem as the evolution of *p*_*ij*_ depends on the joint probabilities *p*(*n*_*ij*_, *n*_*kl*_). The dynamics of the latter depend on the joint occupation probabilities at three or more sites leading to an infinite hierarchy of equations. Following a common approach [28, 29], we assume that the occupation states at neighboring sites are independent, that is, *p*(*n*_*ij*_ = 1, *n*_*kl*_ = 1) = *p*_*ij*_*p*_*kl*_, yielding a closed system of ODEs for *p*_*ij*_ (see Eq. (S2)). The evolution of Eq. (2) and its approximation are both consistent with direct SMM simulations: When *k* > *k*\* we observe an allvertical state (*p*_*ij*_ ≈ 1) when Γ > 1, and an all-horizontal state (*p*_*ij*_ ≈ 0) when Γ < 1. When Γ = 1 orientations tend to be perpendicular to the closer boundary, and *p*_*ij*_ ≈ 0.5 along the diagonals of the square lattice. In Fig. 3c we show the steady-state distribution of cell orientations when *k* > *k*\* and *k* < *k*\* for Γ = 1 (See Fig. S2 for equilibria at different parameter values).

As in the SMM, equilibrium stability depends on Γ and *κ*. Fig. 3a shows the largest real parts of the eigenvalues of the Jacobian of the closed ME at equilibria *p*_*ij*_ = 1 and *p*_*ij*_ = 0 for fixed *k* > *k*\* as a function of Γ. For Γ > 1, the all-vertical state is stable. As Γ crosses unity from above, the largest eigenvalue becomes positive, and the all-vertical state becomes unstable. The all-horizontal state exhibits the opposite behavior. For smaller lattices a saddle-like state (See Fig. 3c and Fig. S2) is stable over a range of Γ (inset in Fig. 3a). Although discrete, Γ thus behaves as a parameter for a transcritical bifurcation in which the all-vertical and all-horizontal states exchange stability with a saddle state.

Consistent with the SMM, when *k* < *k*\*, the equilibria in the regimes Γ < 1, and Γ > 1 are opposite those when *k* > *k*\* (See Fig. S2). Hence, *κ* acts as a second bifurcation parameter for the ME with the all-horizontal and all-vertical equilibria exchanging stability at critical value *k*\*: When Γ > 1, and *k* < *k*\* the stable equilibrium is *predominantly* horizontal. As κ grows, this equilibrium transitions to being *predominantly* vertical, and for some *k* > *k*\*, it destabilizes and the stable equilibrium becomes all-vertical (see Figs. 3b and 4). For brevity, we refer to equilibria only as all-horizontal or all-vertical.

**FIG. 4.**
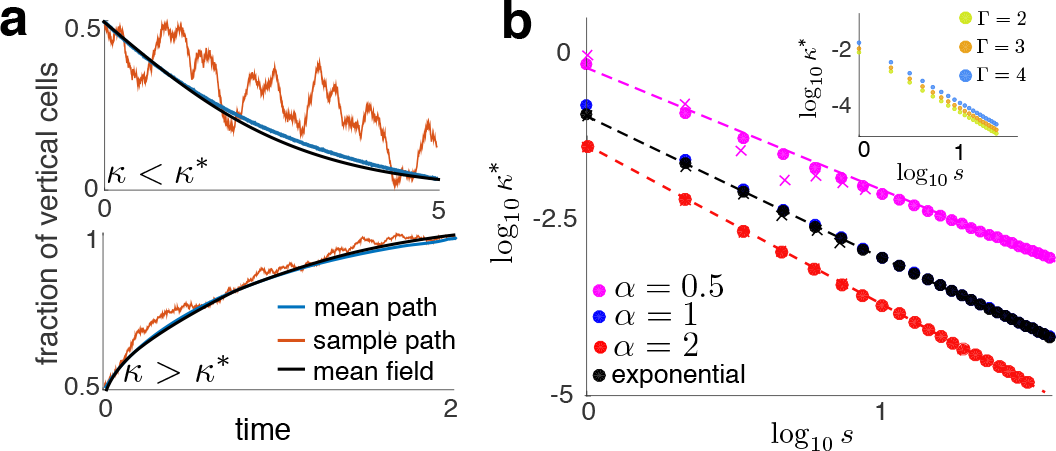
(Color online) (a) Comparison of MF solutions with averages over realizations of the SMM (*N* = 20, *M* = 10). (b) *k*\* as a function of s for different interaction kernels. Dots represent *k*\* values from Eq. (3). X’s were obtained numerically from simulations of the SMM using bisection. Dashed line were obtained using Eq. (4). Inset: *k*\* as a function of *s* for different aspect ratios, Γ.

The transition in stability near *k* = *k*\* and Γ = 1 is driven by the same mechanisms as in the SMM: At Γ = 1 the aspect ratio of the trap changes, while for *k* > *k*\* location-dependent dampening of growth overcomes the loss of cells across the longer trap boundary.

Interestingly, solutions exhibit boundary layers for *k* < *k*\* (see Fig. S2). This suggests a breakdown in the closed ME near the trap’s edges. Indeed, Monte Carlo simulations of the SMM show high correlations between adjacent states near the short trap edge when *k* < *k*\*. However, these correlations decay rapidly away from the boundaries (see Fig. S3).

## Mean field reduction

We next derive a simple mean field (MF) model that captures the behavior of the SMM, and allows us to compute *k*\* analytically. To do so we average occupation states over the lattice. Let

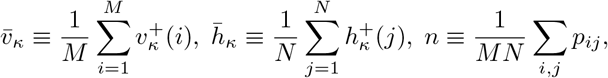

so that *n*(*t*) is the fraction of vertical cells at time *t*, and *v*̄_*k*_, *h*̄_*k*_ are the average growth rates in the vertical, and horizontal directions, respectively. By symmetry, 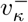, and 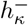 also average to *v*̄_*k*_, and *h*̄_*k*_. Averaging the closed ME over all *i*,*j* shows that *n* obeys a logistic equation,

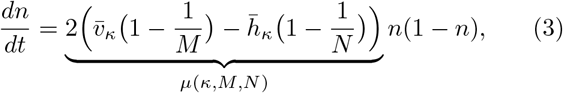

and *n*(*t*) = exp(*μ*(*κ*, *M*, *N*)*t*)/(1+exp(*μ*(*κ*, *M*, *N*)*t*). This agrees with the averaged solutions to Eq. (2), and SMM simulations averaged over realizations (see Fig. 4a).

The effective growth rate of the vertical cell fraction is thus *μ*(*κ*,*M*,*N*). When *κ* = 0, *μ*(0,*M*,*N*) = 2λ(1/*N* − 1/*M*), and the effective growth rate is completely determined by boundary lengths. Cell-cell interactions modulate the effective growth rate as *κ* is increased. However, the system always has two equilibria corresponding to an all-vertical (*n* = 1) and all-horizontal (*n* = 0) orientation which exchange stability at *N* = *M*(Γ = 1).

The two equilibria also exchange stability at a critical level of cell-cell interactions, *k*\*. For fixed *M*, *N*, this transition point satisfies *μ*(*k*\*, *M*, *N*) = 0. For 0 < *k* < *k*\*, and *N* < *M* (*N* > *M*) the state *n* = 1 (*n* = 0) is stable. When *k* > *k*\* the difference in average growth rates, *v̄*_*k*_, *h̄*_*k*_, dominates boundary effects, and the system reaches the opposite equilibrium. Unlike the ME, the MF model predicts a sharp transition between stable equilibria (see Fig. 3b), and no intermediate stable states. Although information about the underlying bifurcation structure is lost, the predicted equilibria and their stability agree with simulations of the SMM, and, when *k* > *k*\*, with experimental observations.

While a general closed form solution for *k*\* is not available, approximate solutions are obtainable for large domains. This allows us to see how *k*\* scales with trap size for different interaction kernels. To reduce parameter number, we fix *M* and *N*, and use a single parameter, *s*, to determine lattice dimensions as *sM* x *sN*. Expanding *μ*(*κ*, *sM*, *sN*) to second order in *κ*, and solving for *k*\* shows that for exponential kernels,

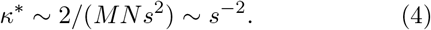

For interaction kernels that decay with the inverse power of distance from the boundary, 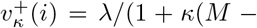
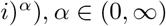,

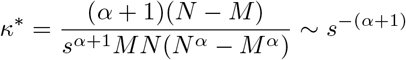

for large s (See SI).

These asymptotic results agree with simulations (See Fig. 4b): *k*\* → 0 as *s* → ∞ at the predicted asymptotic rate. Interestingly, the exponential interaction kernel does not produce the strongest decay of *k*\* with *s*. The aspect ratio of the trap, Γ, shifts the transition points, but does not change the scaling (See inset of Fig. 4b). In large traps even weak cell-cell interactions can cumulatively dominate boundary effects, and lead to steady-state cell alignments orthogonal to the trap’s long side

## Discussion

We showed that cell-cell interactions and boundary effects drive steady-state alignment of bacterial cells in a model of rectangular microfluidic traps: Cell loss across the trap’s edge drives growth parallel to the long side of the trap, while cell-cell interactions drive orthogonal growth. The full stochastic model is well-approximated by a logistic equation which allowed us to analyze the phase transitions in detail.

Similar SMMs have been used to understand tumor initiation and growth [30–33]. However, these models did not include spatially-dependent growth rates and included different boundary conditions. While some analysis is possible, the behavior of these systems is typically less tractable.

Our model is easily extended: We can allow for stochastic switching of orientation, and include more than two orientations. Experiments in rectangular microfluidic devices show that cell alignment destabilizes near the short boundary. Our model extends to capture these features by allowing cell-cell interactions only within a certain distance. Furthermore, we can model multiple bacterial strains by increasing the number of occupational states at a lattice site. Including dynamical equations that describe cellular communication via quorum sensing molecules would then allow us to examine the interplay between cell distribution, communication and growth that determine bacterial collective dynamics [7, 8, 34, 35].

## ACKNOWLEDGMENTS

We thank P. Bressloff and J. Winkle for helpful comments. This work was supported by NIGMS grant R01GM117138 (BRK, MRB, WO, and KJ) and NSF grant DMS-1662290 (BRK, MRB, and KJ).

